# Genome-centric metagenomics reveals novel electroactive syntrophs in a conductive particle-dependent consortium from coastal sediments

**DOI:** 10.1101/2025.08.12.669901

**Authors:** Danijel Jovicic, Konstantinos Anestis, Jacek Fiutowski, Bo Barker Jørgensen, Kasper Urup Kjeldsen, Amelia-Elena Rotaru

**Affiliations:** Department of Biology, Faculty of Natural Science, University of Southern Denmark, Odense, DK; Mads Clausen Institute, SDU NanoSYD, University of Southern Denmark, Sonderborg, DK; Section for Microbiology, Department of Biology, Aarhus University, Aarhus, DK

**Keywords:** syntrophic acetate oxidation (SAO), direct interspecies electron transfer (DIET), conductive particle mediated electron transfer (CIET), methanogens, *Geobacter*, *Methanosarcina*, extracellular electron transfer (EET), *Geobacter psychrophilus*

## Abstract

Conductive particles are abundant in coastal sediments, yet the organisms and pathways that use them for methane production remain unclear. We applied long-read, genome-resolved metagenomics to a sediment-derived consortium that remained dependent on granular activated carbon (GAC) for a decade. We identified a particle-obligate food web of electrogenic syntrophic acetate oxidizers (SAO), an electrotrophic methanogen, and necromass recyclers. The dominant SAO electrogen was a new genus, *Candidatus* Geosyntrophus acetoxidans (<70% ANI/AAI to described taxa), encoding a streamlined extracellular electron-transfer system: one porin-cytochrome conduit (PCC), 47 multiheme cytochromes, conductive pili, and acetate uptake/utilization genes. A second SAO electrogen, *Lentimicrobium* sp., carried two giant PCC-like clusters, suggesting an alternative acetate-oxidation route. Electrons flowed via GAC to a *Methanosarcina* (<89% ANI/AAI to described taxa) equipped with the multiheme cytochrome MmcA and a Rnf/Fpo/HdrDE circuit for EET-driven CO2-reducing methanogenesis. Particle-free lines lost both partners and methanogenic activity, establishing particles as the determinant of persistence. This first genomic blueprint of a natural CIET-SAO consortium identifies potential genomic markers (distinct PCCs, MmcA) for *in-situ* detection and reveals a particle-bound route from acetate to methane likely operating as a fundamental electron-transfer unit in geoconductor-rich anoxic sediments.

## Introduction

Methane is a potent greenhouse gas with a global warming potential approximately 84-times higher than CO_2_ over a 20-year period^1,2^. A considerable fraction of the atmospheric methane is derived from acetate, channeled either through acetoclastic methanogens (Eq. 1) or by syntrophic acetate oxidation (SAO), in which an acetate-oxidizing bacterium supplies reducing equivalents (e.g., H_2_ or electrons) to a partner methanogen, rendering acetate oxidation thermodynamically feasible^3,4^. (Eq. 2-3). Although these reactions operate at the thermodynamic limits of life, SAO is essential in methane-producing ecosystems.

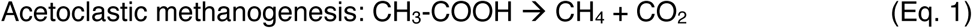

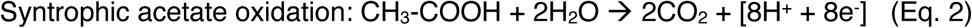

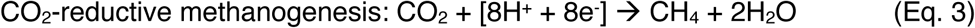

SAO-driven methane formation occurs across different anoxic environments including anaerobic digesters^5–8^, rice paddies^9–11^ and freshwater or marine sediments^12–15^. In coastal settings, natural and climate driven erosion mobilizes terrigenous material^16^, such as iron oxides (e.g., magnetite)^17,18^ and wildfire-derived char particles ^19–21^ that act as semiconductor conduits^22^. Laboratory studies show that such particles or their analogues (i.e., granular activated carbon, an analogue for chars) can raise methane production up to 15-fold by serving as electrical conduits for CIET between electrogenic bacteria and electrotrophic methanogens in synthetic consortia^23–27^. A similar, four-fold stimulation of methane production was obtained when supplying conductive particles from onset to laboratory enrichments from aquatic sediments (coastal marine or lacustrine)^15,22^, without the particles the native CIET-SAO consortia were lost. These particle-dependent partnerships may determine whether acetate (a central intermediate in conversion of organics to methane) is routed to methane via SAO or directly via acetoclastic methanogenesis. Because CIET-SAO remains virtually unreported *in situ*, and its mechanisms remain elusive, these sediment enrichments initiated with conductive particles remain the closest practicable proxy for disentangling how these consortia route acetate to methane and accelerate organic-matter turnover in anoxic soils and sediments^4^.

The first compelling evidence for conductive particle-obligate SAO came from enrichments established from coastal Bothnian Bay sediments^15^. When singly methyl-labelled acetate (^13^CH_3_-COOH) was supplied, ^13^CO_2_ accumulated only in GAC-amended bottles, and never in GAC-free controls, indicating acetate oxidation by bacteria rather than acetoclastic cleavage by methanogens. NanoSIMS confirmed that more than 80% of the ^13^C-label was indeed incorporated in a *Geobacter-*like bacterium, partnered with a *Methanosarcina-*like archaeon. Yet 16S-phylogeny and probes alone could not resolve the phylogeny of either organism (Suppl. Fig. 4-5). The *Geobacter*-like bacterium shared only 95-96.6% identity to *Geobacter psychrophilus,* a taxon peripheral to canonical *Geobacteraceae*. The *Methanosarcina* matched 98-99% to both *M. subterranea* and *M. lacustris*. Inhibitor tests showed that blocking methanogenesis immediately stopped acetate oxidation, confirming strict co-dependency between partners. Remarkably the consortium proved GAC obligate: without GAC, methane production dropped four-fold, and the CIET-SAO partners declined to undetectable levels, allowing a strict acetoclastic methanogen to dominate^15^. Despite this compelling physiological evidence, the phylotypes, electron-transfer routes, and genomic determinants that enable these novel electrogenic and electrotrophic partners to thrive on conductive particles remained unresolved.

In this work we closed this gap by applying long-read, genome-centric metagenomics to the above Bothnian Bay consortium serially propagated for a decade on GAC. We recovered 24 metagenome assembled genomes (MAGs), including near-complete genomes of the SAO *Geobacter*-like bacterium and its *Methanosarcina* partner alongside other coexisting taxa. We could identify the conductive-particle-linked electron-transfer machinery and reconstruct the metabolic network that channels acetate to methane. These results provide the first genomic blueprint of a conductive particle-obligate SAO community and a foundation for assessing how natural and anthropogenic conductive particles could amplify methane emissions in coastal marine environments.

## Results and Discussion

After a decade of serial transfers with GAC, the Bothnian Bay SAO consortium continued to oxidize acetate at 0.5 ± 0.1 mM day^-1^ and produce 0.2 ± 0.0 mmol CH_4_ day^-1^, matching previously reported rates^15^ (**Fig. 1**a). By contrast, a parallel line of serially transferred GAC-free controls remained virtually inactive over the 42-day incubation period (acetate: 0.01 ± 0.09 mM day^-1^; methane: 0.01 ± 0.02 mmol day^-1^) (**Fig. 1b**), confirming that the electrically conductive scaffold provided by GAC is still indispensable for SAO after a decade of serial cultivation and enrichment.

Helium-ion micrographs of GAC-particles (**Fig. 1c-e**) revealed round *Methanosarcina-*like cells (M) interspersed among rod-shaped bacterial partners (B) without consistent cell-to-cell contact, supporting the notion that electrons are exchanged via the conductive matrix rather than through direct cell-to-cell contact.

**Figure 1.**
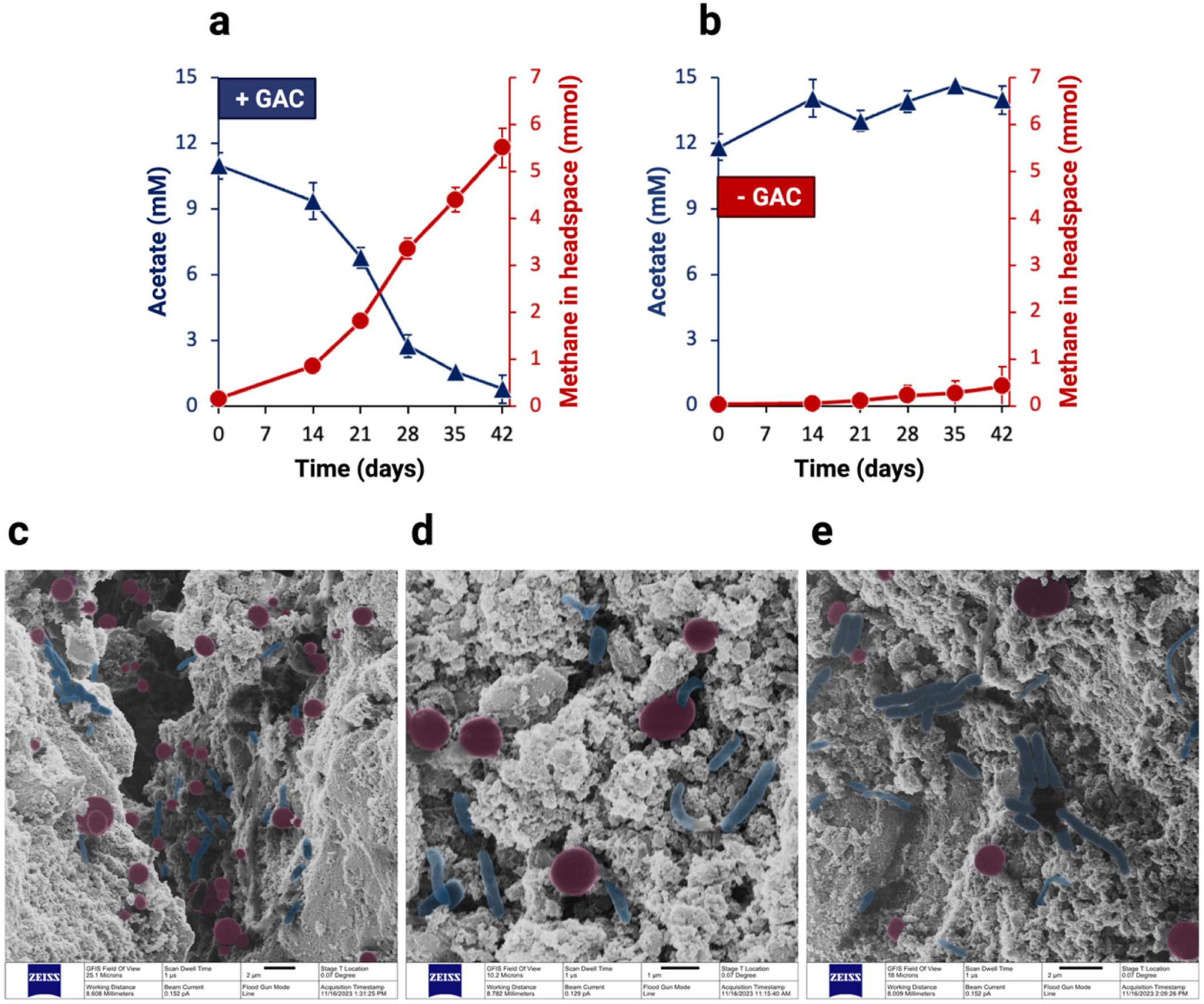
Syntrophic acetate oxidation (SAO) by a Baltic Sea consortium dependent on conductive particles. (a, b) Methane production (●) and acetate consumption (▲) over 42 days in anaerobic cultures incubated (a) with granular activated carbon (GAC) or (b) without GAC; values are means ± SD of biological replicates (n ≥ 3). (c–e) Colorized helium-ion microscopy (HIM) micrographs of GAC particles at day 42 showing *Methanosarcina* cells (red, round cells) and associated bacterial partners (blue, long and short rods). Scale bars: 2 μm (c, e) and 1 μm (d). Additional HIM micrographs are included in the Supplementary Material (Suppl. Fig. 1-2).

To assess the identity of the SAO-bacteria and partner methanogen that continued to inhabit GAC-particles after a decade of sediment free serial propagation, we performed metagenomic profiling of the GAC-obligate microbiome. Our analysis revealed indeed a MAG far related to *Geobacter* accounting for 9.4% of the total reads, along with a *Methanosarcina-*related MAG that represented 12% of the total reads. The 16S rRNA-gene of these two MAGs matched (∼99% and above) the previously reported 16S-rRNA phylotypes^15^. Both these MAGs were essentially absent from GAC-free controls (< 0.1%; **Fig. 2a**). These results are consistent with our earlier stable isotope labelling studies that identified a *Geobacter*-like organism as the dominant acetate oxidizer (incorporating >80% of the labelled acetate) and a *Methanosarcina* as the sole methanogen in the consortium^15^.

In this study, we reconstructed high-quality MAGs (>97% completeness, <0.7% contamination) for the key syntrophic partners, *Geobacter-*like and *Methanosarcina*-like microorganisms (**Fig. 2a**). Beyond these core SAO partners, GAC also enriched specifically the relative abundance of three other species – belonging to *Sedimentibacter* (6.5%), *Lentimicrobium* (3.4%), and *Desulforhopalus* (3%) (**Fig. 2a**). Given that other representatives of *Sedimentibacter* and *Lentimicrobium* were detected in the metagenomic libraries of both GAC-supplemented and GAC-free controls, we hypothesize that their primary ecological role may not be related to direct electron transfer but rather to the recycling of organic matter resulting from biomass turnover within the community.

Taken together, the activity profiles, spatial organisation and metagenomic data indicate that decade long serial transfers of the consortium had not altered its core partnership between *Geobacter*-like bacteria and *Methanosarcina*-like archaea. So, the Baltic Sea *Geobacter* continued to oxidize acetate and transfer electrons through the GAC matrix, while the Baltic *Methanosarcina* completed the syntrophic process by accepting the electrons from GAC to reduce CO₂ to CH₄.

**Figure 2.**
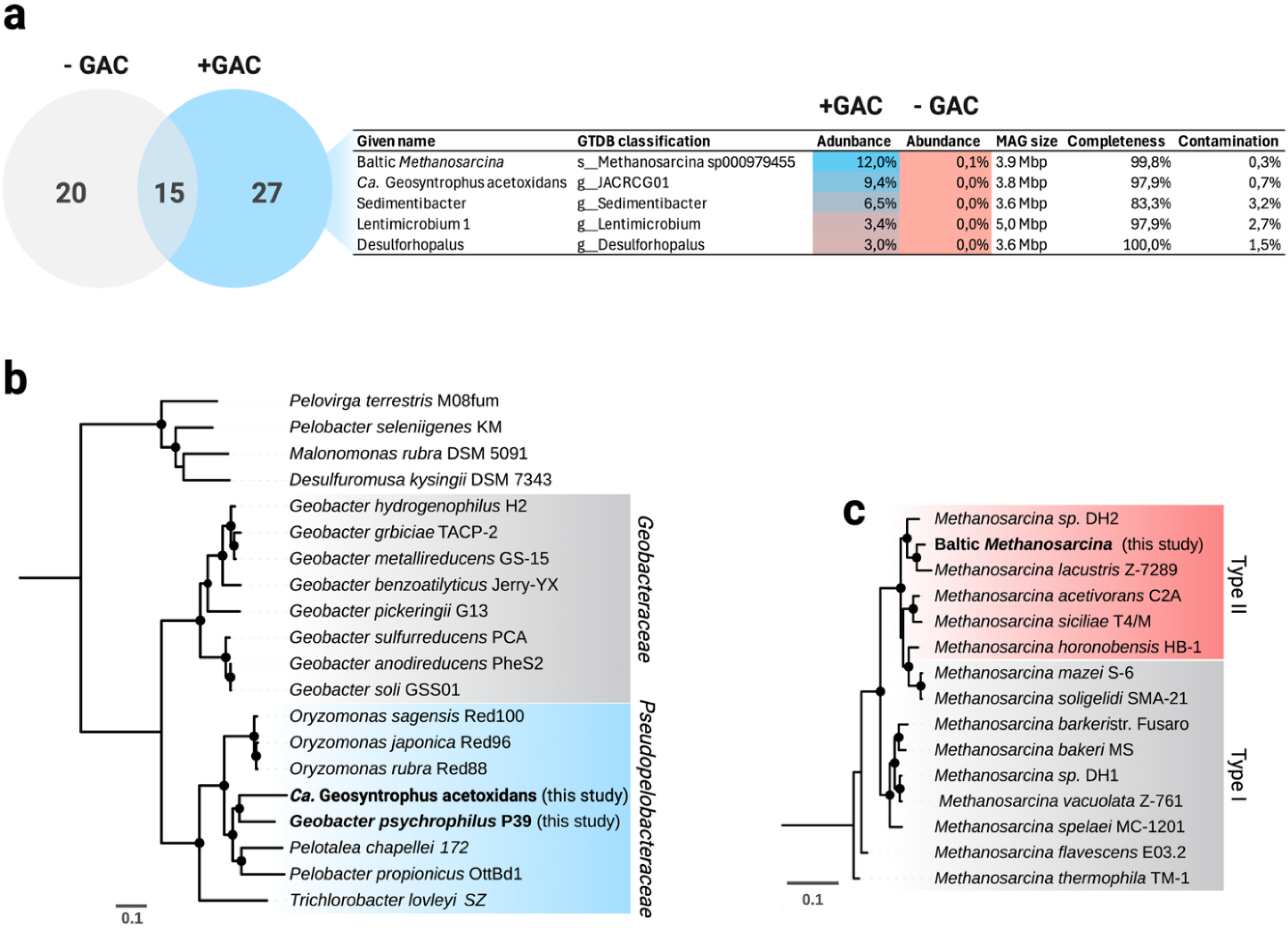
Phylogenomic placement of the key players in a Baltic Sea syntrophic acetate oxidizing consortium dependent on GAC. (a) Venn diagram showing MAGs unique to particle-free controls (−GAC, grey), unique to GAC-amended cultures (+GAC, blue) or shared between both (overlap). The table lists the five most abundant MAGs in the GAC-amended incubations with their GTDB affiliation, relative abundance, genome size, and quality metrics. (b) Maximum-likelihood phylogenomic tree based on a concatenated alignment of 120 GTDB marker proteins. The tree was inferred using RAxML-HPC2 with the PROTCAT+DAYHOFF model and 100 bootstrap replicates. The Baltic electrogen previously listed as a Baltic ‘Geobacter’ here re-named *Ca.* Geosyntrophus acetoxidans (bold) and Geobacter psychrophilus P39 (genome sequenced in this study, bold italics) form a clade within the family Pseudopelobacteraceae (blue shading), clearly delineated from Geobacteraceae (grey shading). per site. **(c)** Corresponding archaeal tree based on 53 archaeal marker genes. The Baltic Methanosarcina (bold) groups with type II strains (red shading) and is distinct from type I lineages (grey shading). For b and c, filled circles mark bootstrap support above 70 % for 100 replicates. Scale bar, 0.1 substitutions per site. Distribution of the MAGs identified under GAC and particle-free incubations can be found in Suppl. Fig. 3.

### The *Geobacter*-like SAO syntroph represents a novel genus, *Ca.* Geosyntrophus acetoxidans

Despite its stable dominance over a decade of serial transfers, the phylogenetic identity of this *Geobacter*-like bacterium remained unresolved. A 16S-rRNA gene phylogeny yielded ambiguous results, placing the Baltic ‘*Geobacter’* between *Pelobacter chapellei* and *Geobacter psychrophilus* (sharing ∼97% identity with both, Suppl. Fig. 4). Furthermore, the Genome Taxonomy Database (GTDB) classified the ‘*Geobacter’*-MAG into an uncharacterized genus level lineage (genus JACRCG01).

Because no genome information existed for its closest relative, *Geobacter psychrophilus*, we sequenced the genome of the only available strain of this species – *G. psychrophilus* P39 (Nevin et al., 2005), kindly provided by Prof. D. Lovley (UMass Amherst). Concatenated phylogenomic analysis using 120 marker genes^28^ placed the Baltic ‘*Geobacter’* firmly in a robust clade with *G. psychrophilus* P39. Yet their genomic divergence was substantial, sharing only 73.3% average nucleotide identity (ANI) and 68.8% average amino acid identity (AAI) values that fall at the limit, or below the limit, of accepted genus thresholds (∼75 % ANI, 60-80 % AAI, Suppl. Fig. 6)^29,30^.

Consistent with a comprehensive reevaluation of the order *Geobacterales* by Xu et al. (2021)^29^, the Baltic Sea ‘*Geobacter’* and *G. psychrophilus* clustered within a newly proposed family *Pseudopelobacteraceae*^31^, clearly separated from canonical *Geobacter* species of the family *Geobacteraceae* (**Fig. 2b**). This taxonomic ambiguity of the order *Geobacterales*, emphasizes the urgent need for a formal revision. To clearly delineate the Baltic *’Geobacter’* from other *Pseudopelobacteraceae* we propose the name *Ca*. Geosyntrophus acetoxidans, marking its novelty and ecological specialization.

High quality genome reconstruction (97.9% completeness and 0.7% contamination) of *Ca*. Geosyntrophus acetoxidans MAG encodes the entire toolkit for acetate oxidation that is seamlessly coupled to an all-inclusive extracellular electron-transfer (EET) apparatus including conductive pili and 47 multiheme cytochromes (MHCs), the largest MHC repertoire among all the MAGs recovered (**Fig. 3**). The EET apparatus of *Ca*. Geosyntrophus had similar modules and architecture to that of *Geobacter* species, however neither of the constituents showed significant sequence identity to any canonical *Geobacter*.

Acetate enters the cell cytoplasm through porins and an ActP a cation/acetate symporter (*actP*) and it’s activated to acetyl-CoA either directly via acetyl-CoA synthetase (*acs*) or with intermediate formation of acetyl-phosphate catalyzed by acetate kinase (*ackA*) and phosphate acetyltransferase (*pta*). The resulting acetyl-CoA is then completely oxidized via a complete tricarboxylic acid cycle (TCA), generating NADH and FADH_2_. NADH feeds electrons to the membrane-bound NADH-quinone oxidoreductase (*nuoA-N*) reaching the menaquinone pool and driving proton translocation. FADH_2_ generated by succinate dehydrogenase funnels also electrons from TCA to the menaquinone pool. From menaquinol, electrons could pass to nine inner-membrane MHCs — including ImcH/CbcL homologues—and enter a densely populated periplasm containing 28 MHCs (Supplementary Table 1). Two of these share low (∼45%) amino acid identity with well described PpcA/PpcD redox hubs of *Geobacter* spp.^32,33^, reflecting functional convergence despite distant phylogeny.

Periplasmic electrons exit through a single porin-cytochrome-cytochrome (PCC) conduit (OmbB-OmaB-OmcB). PCC conduits have been shown to be required for EET, and in the absence of all PCCs extracellular electron transfer is abolished in *Geobacter*^34^. In *Ca*. Geosyntrophus, OmaB (8 hemes) and OmcB (12 hemes) mirror the heme architecture of the archetypal PCC of *G. sulfurreducens*, but they share less than 30 % amino-acid identity, indicating independent evolution of an identical solution. The presence of only one PCC cluster—versus multiple sets in most *Geobacter* spp.—is consistent with our consortium’s strict reliance on the conductive particle, but inability to pair with the methanogen directly.

**Figure 3.**
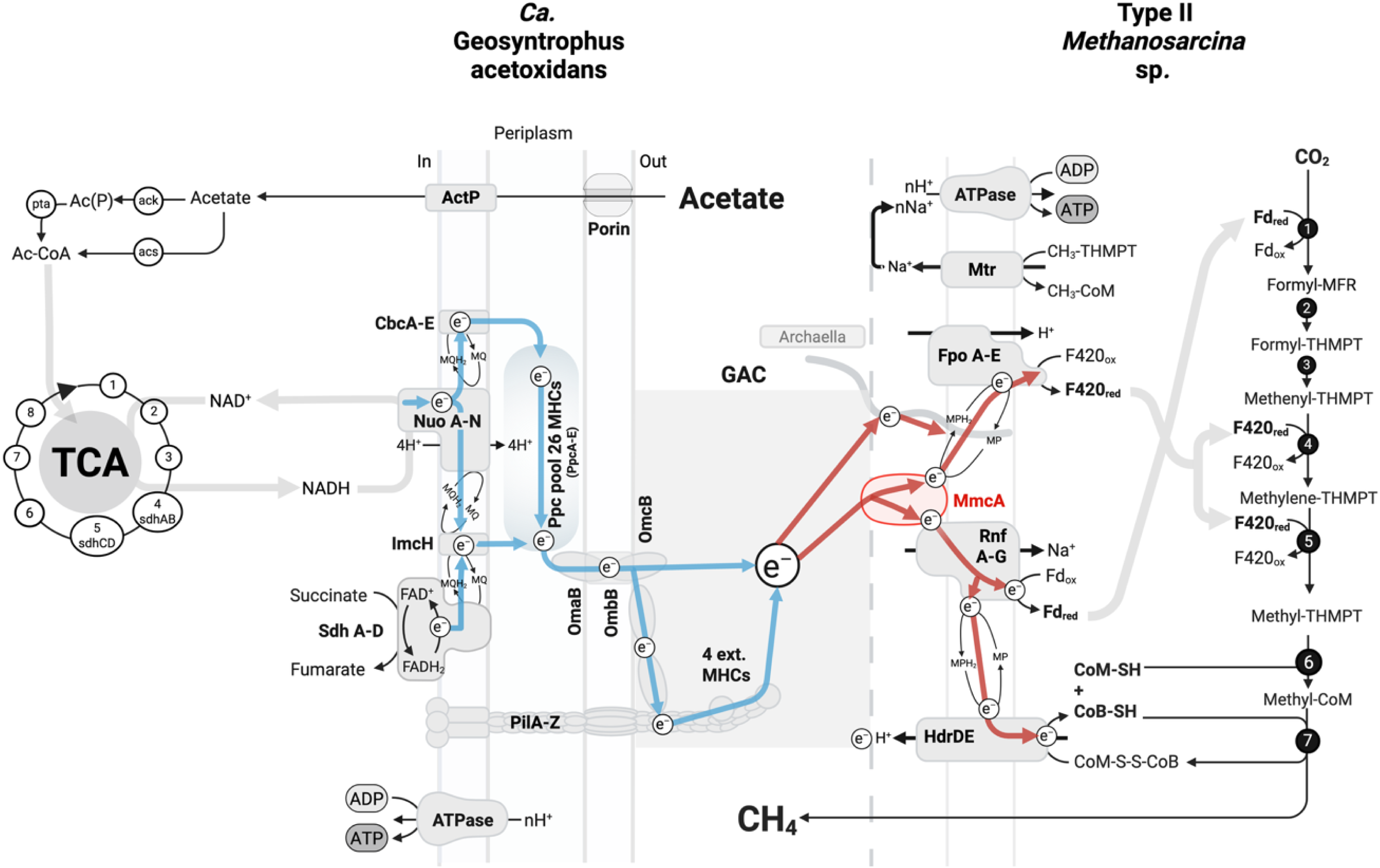
Proposed electron-transfer network for syntrophic acetate oxidation using GAC. The metabolic reconstruction incorporates high-quality MAGs of *Ca.* Geosyntrophus acetoxidans (left, blue) and Baltic type II Methanosarcina sp. (right, red). In *Ca.* Geosyntrophus acetoxidans, acetate is imported and activated to acetyl-CoA, which is fully oxidized in the TCA cycle, producing NADH and FADH₂. Electrons from these cofactors enter the menaquinone pool via NADH dehydrogenase (Nuo) and succinate dehydrogenase (Sdh), flowing through inner-membrane MHCs to a periplasmic reservoir before exiting the cell via a PCC-conduit. Extracellular MHCs and conductive e-pili transfer electrons to GAC, which then reach the methanogen. Electrons from GAC arrive at the outer-surface of the methanogen to reach the MHC MmcA or, secondarily, at an archaellum-associated route (grey wire-like). MmcA contributes electrons to reducing the methanophenazine (MPH_2_) pool, bifurcating electrons to the Rnf complex and F₄₂₀H₂ dehydrogenase. Besides ferredoxin, Rnf also supplies electrons to heterodisulfide reductase, regenerating reduced Coenzyme M and Coenzyme B. The TCA-cycle and the CO_2_-reductive methanogenesis pathway involve several enzymes, all encoded in the MAGs of *Ca.* Geosyntrophus acetoxidans and Baltic Methanosarcina, respectively. Reactions 1-7 for CO_2_-reductive methanogenesis: (1) CO_2_-activation by methanofuran, (2) formyl transfer to tetrahydromethanopterin (H_4_MPT), (3) cyclization to methenyl-H_4_MPT, (4) reduction to methylene-H_4_MPT, (5) further reduction to methyl-H_4_MPT, (6) methyl transfer to coenzyme M, (7) final reduction of methyl-coenzyme M to methane.

Beyond the outer membrane, electrons may be relayed either by extracellular MHCs that can polymerise into nanowires and/or by two PilA monomers whose aromatic-residue pattern is identical to that of the conductive pili of *G. metallireducens* (11.5 % aromatics; **Fig. 4**). Previously, it was shown that the aromatic amino acid distribution is essential for electron delocalization and long-range conductivity^35,36^. Also, deletion of *pilA* abolished DIET between *Geobacter* and a type II M*ethanosarcina* (*M. acetivorans*) and GAC could completely compensate for this loss suggesting that the role of pili in the interaction with GAC and the methanogenic partner is not crucial^37^.

Taken together, these data indicate that *Ca.* Geosyntrophus acetoxidans oxidizes acetate and disposes of the resulting electrons through a streamlined, particle-obligate EET-conduit. Electrons from the MHC-rich periplasm are exported through a single porin-cytochrome conduit and then relayed by four extracellular MHCs and the pili to the conductive GAC matrix. Unlike canonical *Geobacter* species, which encode multiple PCC sets to engage diverse electron acceptors or partner cells in a changing environment^38–40^, *Ca*. G. acetoxidans retains only one. This minimal redundancy is sufficient for CIET, but limits alternative strategies such as direct cell-to-cell exchange^15^. Originating from the conductive-particle-rich, relatively stable deeply anoxic sediments of the Bothnian Bay, *C*. Geosyntrophus likely shed redundant EET pathways as an adaptation to its niche. These features also exemplify convergent evolution of the *Geobacter*-type EET conduit outside the *Geobacteraceae* family.

**Figure 4.**
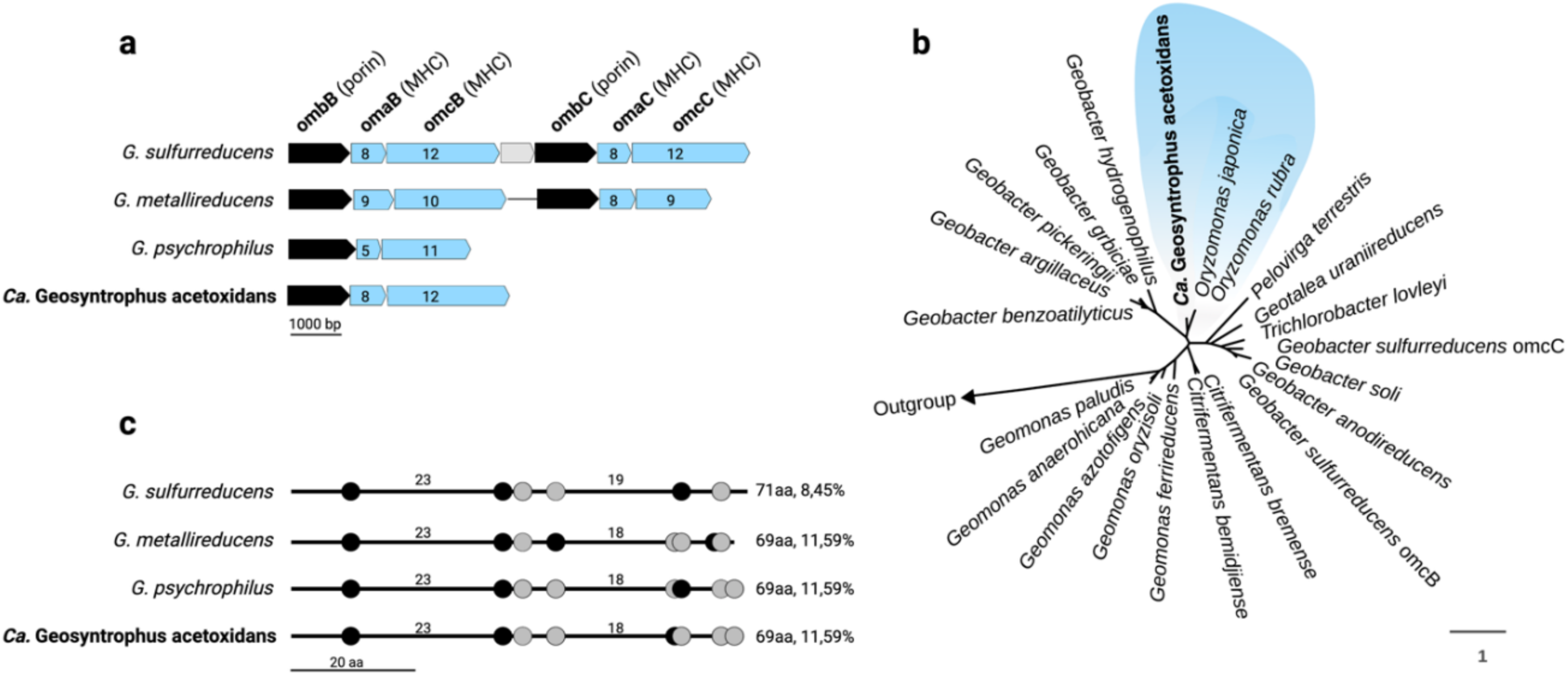
Outer-surface EET conduit of *Ca.* Geosyntrophus acetoxidans compared with its closest relative *G. psychrophilus* P39 and benchmark Geobacter species. (a) Representation of Porin-Cytochrome-Cytochrome (PCC) gene clusters in *Ca*. Geosyntrophus acetoxidans versus other Geobacter, detailing the number and arrangement of heme-binding motifs. (b) Maximum likelihood phylogenetic tree of the OmcB aminoacid sequence from representative *Geobacterales*. The tree was inferred using RAxML-HPC2 on the CIPRES Science Gateway, based on protein alignments (1323 aligned amino acids) using the PROTCAT+DAYHOFF model with 100 bootstrap replicates. The OmcB of *Ca.* G. acetoxidans (blue shading) forms a well-supported clade with other *Pseudopelobacteraceae* (*Oryzomonas*) and is clearly separate from the *Geobacteraceae* core. Scale bar represents 1 amino acid substitution per site. (c) Linear map of aromatic residues along PilA monomers that form conductive e-pili annotated with the positions of aromatic amino acids: tyrosine black (Y) and phenylalanine grey (F). The *Ca*. Geosyntrophus acetoxidans pilA replicates the spacing found in the highly conductive PilA of *G. metallireducens*; values on the right give total length (aa) and percentage of aromatic residues. Scale bar, 20 aa. Additional information about multiheme cytochrome content per MAG can be found in Suppl. Fig. 8. Additional comparison of pili amino acid sequences and aromatic amino acid content between *Ca*. Geosyntrophus and 18 other species can be found in Suppl. Fig. 9.

### Methanogenic partner is a novel type II *Methanosarcina* species

Phylogenomic analysis of 53 concatenated archaeal marker genes place the Baltic Sea *Methanosarcina* MAG within the Type II (non-H_2_-utilizing) *Methanosarcina* clade^41^, next to *M. lacustris* and *Methanosarcina sp.* DH2 (**Fig. 2c**). Whole genome comparison shows only 89% ANI (Suppl. Fig. 7) and 85.2% AAI to its nearest relative, *M. lacustris* – well below the ∼95% ANI species boundary, supporting that it represents a novel Type II *Methanosarcina* species.

The Baltic Sea *Methanosarcina* MAG is near-complete (99.8 % completeness, <0.3 % contamination) and encodes the full complement of genes for acetoclastic, methylotrophic and CO₂-reducing methanogenesis, but—characteristic to type II Methanosarcina clade— it lacks uptake Ech-hydrogenases^41^. In terms of bioenergetics, this *Methanosarcina* has a complete set of genes for the sodium-pumping Rnf complex (*rnfABCDEG*) together with the membrane-bound F₄₂₀H₂ dehydrogenase (*fpoABCDHIJKLMN*) and heterodisulphide reductase (*hdrDE*), providing the canonical Rnf/Fpo circuit that drives ATP synthesis during acetate or CO₂ reduction coupled with DIET or CIET^42^ (Supplementary Table 1). Crucially, the genome encodes a multi-heme c-type cytochrome MmcA. The MmcA is a well characterized bidirectional redox conduit shown to be indispensable for direct interspecies electron transfer (DIET) and Fe(0) oxidation in a type II *Methanosarcina*, *M. acetivorans*. A deletion of *mmcA* abolishes both electron uptake and electron transfer in this methanogen^37,43–45^.

In our previous work we have shown that this same Baltic Sea *Methanosarcina* (≥99% 16S rRNA sequence identity) was incapable of H_2_ or acetate utilization^46^, but it reduces CO₂ efficiently when electrons are supplied by Fe(0)^46^ or *Ca. Geosyntrophus acetoxidans* via GAC^15^. Methane production collapses if the electrogenic partner or conductive particles are removed, suggesting that this methanogen is obligately reliant on extracellular electron uptake.

We recovered the MAG of this methanogen and show that it contains the full genetic make up for extracellular electron uptake and CO_2_ reductive methanogenesis (**Fig. 3**). We propose the following working model: each four (of the eight) electrons liberated from acetate oxidation by *Ca*. Geosyntrophus acetoxidans are delivered to GAC, accepted by the outer-surface MHC, MmcA, and then bifurcated. Two electrons reach the Rnf complex generating reduced ferredoxin; the other two enter the methanophenazine pool to fuel a reverse-running Fpo that reduces F_420_ while importing protons. Although reversal of the Fpo consumes the proton gradient, this investment is counterbalanced later in the pathway: the Na⁺-pumping methyltransferase Mtr exports Na⁺ during the methyl-H₄MPT→CoM step, and membrane HdrDE reduces CoM-S-S-CoB via the methanophenazine pool with scalar H⁺ release to the periplasm-like space, rebuilding the ion motive force^42^ (**Fig. 3**). Thus, the extracellular electrons reduce all carriers (ferredoxin and F_420_) necessary to initiate CO₂ reduction to methane, while Rnf simultaneously supplies electrons to HdrDE to generate reduced CoM and CoB for the terminal step of the CO_2_-reductive methanogenesis pathway. The regenerated ΔμH⁺/ΔμNa⁺ then drives ATP synthase, which can use either ion in *Methanosarcina* ^42^.

The genome also encodes two copies of a complete archaellum operon, including two flagellin genes (>60% amino acid similarity to *M. acetivorans* FlaB1 and FlaB2), that provide a potential alternative path for extracellular electrons. However, GAC fully compensated for the loss of archaellin in a Δ*flaB1*Δ*flaB2* mutant of *M. acetivorans*^37^, indicating that the archaella becomes dispensable when long-range electron transfer can proceed via conductive particles like GAC.

The electron-uptake machinery (MmcA and archaellin) in Baltic Methanosarcina parallels the mechanism experimentally resolved in DIET-grown *M. acetivorans*^37^, suggesting that environmental type II Methanosarcina come equipped with a comparable genetic toolkit to perform EET.

While direct-electron partnerships (DIET or CIET) involved in methane production have so far been documented mainly in synthetic co-cultures assembled in the laboratory, the Baltic Sea consortium represents a naturally occurring syntrophic partnership. The pairing of a novel electrogenic acetate oxidiser, *Ca. Geosyntrophus acetoxidans*, and a novel Type II *Methanosarcina* sp. introduces two previously unrecognised partners into the catalogue of conductive particle syntrophies. Their reliance on conductive particles, rather than direct cell-to-cell contact, highlights an ecological strategy in which co-occurring respiratory bacteria and methanogens exploit environmentally available geoconductors to exchange electrons, a mechanism that is probably widespread yet largely overlooked in energy-limited sediments.

**Figure 5.**
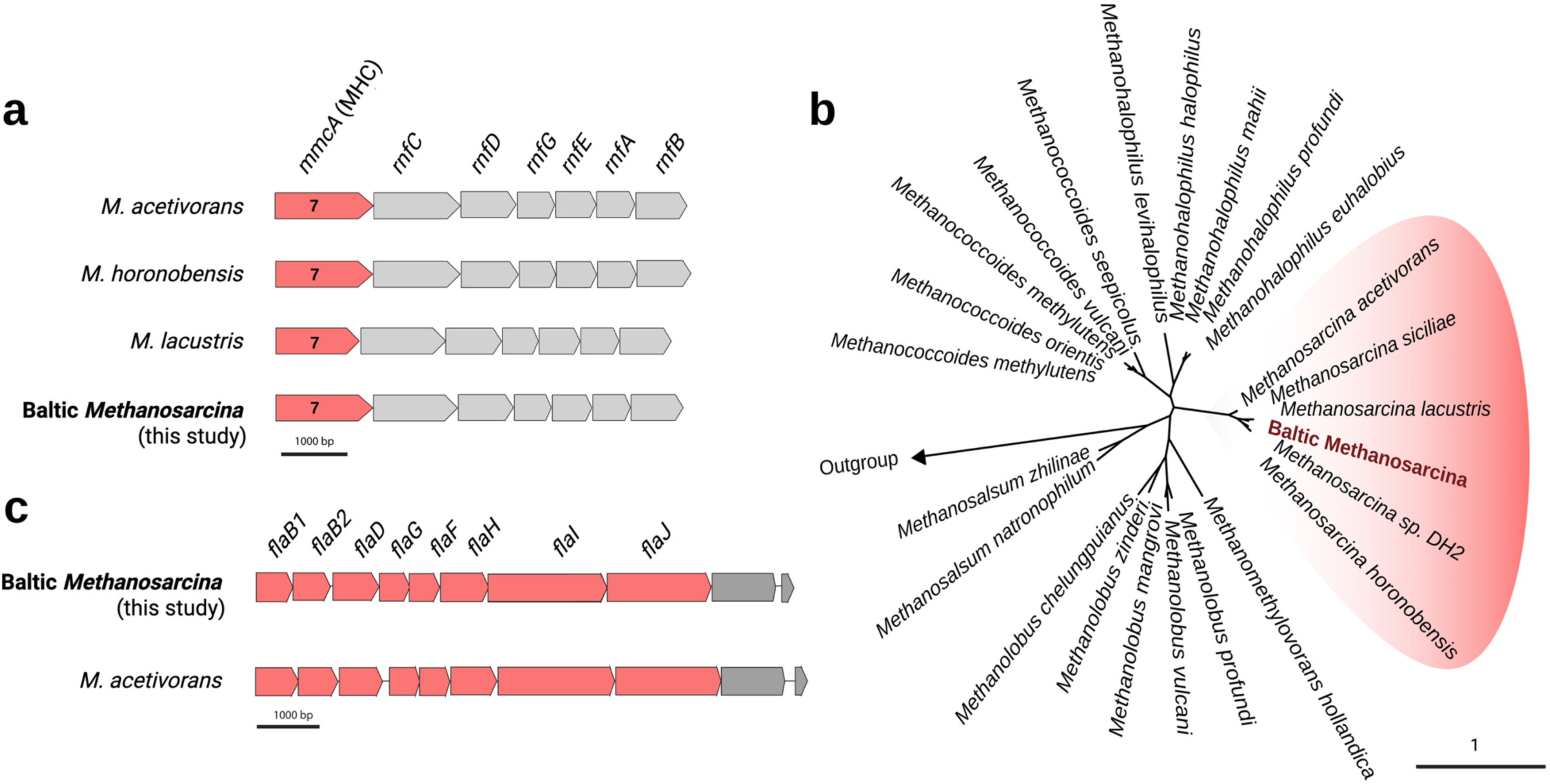
Electron-uptake machinery in the Baltic Sea type II *Methanosarcina*. (a) Gene neighbourhood of the *mmcA*-electron-transfer locus (*mmcA–rnfCDEGAB*) in the Baltic *Methanosarcina* compared to three reference type II strains. Arrows indicate ORF orientation; numbers inside mmcA denote predicted heme-binding motifs. Scale bar, 1 kb. (b) Maximum-likelihood tree of MmcA aminoacid sequences from Baltic Methanosarcina (bold red) clusters with other type II Methanosarcina homologues (red shading) and is clearly separated from MmcA-like cytochromes in other methanogens. Scale bar, one amino-acid substitution per site. (c) Schematic comparison of the complete archaellum operon (archaeal flagellum) in the Baltic genome versus *M. acetivorans*; both operon copies are conserved in gene content and length (grey arrows), supporting the potential for an auxiliary, archaella-based electron-uptake route.

### Auxiliary taxa may engage in biomass turnover or a share of the acetate oxidation

Although *Ca.* Geosyntrophus acetoxidans and the Baltic Type II *Methanosarcina* dominate the GAC-attached consortium, members of three auxiliary lineages, *Sedimentibacter*, a *Desulforhopalus* and a *Lentimicrobium,* were abundant in the GAC-incubations (**Fig. 2a**). Various other *Sedimentibacter* and *Lentimicrobium* MAG-variants were also well represented in particle-free controls (Suppl. Fig. 3), suggesting that these taxa are unlikely to play a role in interactions with the conductive particles.

*Sedimentibacter* (6.4% of the reads in GAC-incubations) are well-known fermenters in methanogenic environments^47–49^ and were present in acetate-fed microbial fuel cells^50^. However, *Sedimentibacter* have, to date, not been implicated in extracellular electron transfer or syntrophic acetate oxidation. We therefore ascribe their enrichment to turnover of cellular detritus released from the core CIET partnership rather than to direct involvement in extracellular electron flow.

*Desulforhopalus* (3% of the reads in GAC-incubations) was unique to GAC incubations. Although species of *Desulforhopalus* can act as incompletely oxidizing sulphate reducers^51^, a sulphate reducing lifestyle is not favoured by the conditions provided by our SAO medium lacking sulphate. Besides, the absence of acetate transporters suggests that it is not involved in acetate uptake for subsequent oxidation. If acting as an incomplete oxidizer, any acetate formed intracellularly can still be released—via passive diffusion of the protonated form (acetic acid) and/or broad-specificity monocarboxylate transporters—so the absence of dedicated acetate importers does not preclude acetate export. In our system, *Desulforhopalus* may be involved organic matter fermentation. Species of *Desulforhopalus* like *D. singaporensis* have been shown to ferment taurine or pyruvate^52^, which provides them with an independent carbon-and energy-acquisition route that does not require direct participation in extracellular electron transfer. Besides, our *Desulforhopalus* MAG lacked PCCs, but contained an unusually long pilA-like gene whose aromatic residue pattern matches that of conductive e-pili, implying a possible role in EET. However, with an incomplete EET machinery and lacking acetate transporters, *Desulforhopalus* is unlikely to be involved in syntrophic acetate oxidation but may be involved in biomass turnover like *Sedimentibacter*.

A single *Lentimicrobium* MAG, representing 3.4% of the GAC community, encoded two large porin–multi-heme cytochrome c clusters (PCC3-like architectures containing 61- and 43-heme modules) and genes for acetate uptake. Members of the family *Lentimicrobiaceae* are characterized by their ability to ferment carbohydrates under anaerobic conditions, producing acetate as a major metabolic by-product^53^. However, previous studies using stable-isotope probing have demonstrated acetate incorporation by *Lentimicrobiaceae* in anaerobic digester communities, suggesting they may act as acetate oxidizers^54,55^. The presence of both PCCs and acetate transporters in this *Lentimicrobium* MAG indicates that Baltic *Lentimicrobium* could oxidise acetate and discharge electrons to the GAC matrix, thereby feeding its syntrophic *Methanosarcina* partner. In our previous work, *Ca.* Geosyntrophus (aka Baltic *Geobacter*) was the primary organism taking up labelled acetate^15^ (∼80% incorporation in mid-exponential^15^), We therefore predict that either *Lentimicrobium* is a slower strategist or has different kinetics of acetate uptake and utilization compared to *Ca*. Geosyntrophus.

In contrast, the particle-free controls —while minimally active (see Fig. 1b)—failed to sustain any electrogen-methanogen partnership. Both the GAC-amended and particle-free lines were initiated a decade ago from the same sediment core slurry, but only the former contained conductive particles and supported the growth of electroactive consortia^15^. In particle-free controls, the community instead comprised of a broader array of fermenters, three *Sedimentibacter*, two *Lentimicrobium*, and assorted *Humidesulfovibrio* (Suppl. Fig. 3)—likely involved in organic matter recycling. The absence of an electrogen–methanogen consortium in particle free controls at the 30-st transfer is consistent with our earlier findings from isotope labelling and CARD FISH coupled with NanoSIMS at the 6-th transfer^15^. In these particle-free lines, abundant fermentative taxa are likely to scavenge lysis products from transient populations unable to establish sustained methanogenic growth. Whole genome sequencing after 30+ successive transfers confirmed their poor methanogenic potential: archaeal populations remained at ≤0.5% abundance with a *Methanomassilicoccales* MAG (UBA472) at 0.5% detected only in particle-free incubations, and the *Methanosarcina* MAG barely detectable (0.06%) (Suppl. Fig. 3). Together these results indicate that without conductive particles neither electrogenic syntrophs nor partner methanogens can persist.

These decade long enrichments show that the presence or absence of conductive particles determines whether specific metabolic partnerships persist or collapse, directly shaping carbon flow and methane output. In our consortium, the electrogenic partners, *Ca.* Geosyntrophus acetoxidans and Lentimicrobium both encode PCC clusters that are unrelated to those of canonical *Geobacter* spp. pointing at a convergent evolution of EET machineries in phylogenetically distant lineages. The dominant SAO-bacterium, *Ca.* Geosyntrophus acetoxidans, carries only a single PCC, unlike the multiple PCC sets typical of *Geobacter*^56^. In *Geobacter,* this redundancy enables versatility and ability to rapidly adjust to changing conditions, for example by employing specialized PCCs matched to different extracellular electron acceptors^39,57^. By contrast, the minimal PCC repertoire of *Ca*. Geosyntrophus suggests adaptation to stable, particle-rich, anoxic niches where reliable conductive scaffold precludes the need for alternative EET routes. Taken together, our study provides the first genomic blueprint of a naturally occurring microbial consortium obligately reliant on conductive particles. By extending conductive particle mediated syntrophy beyond synthetic dual-species laboratory models, we could begin to grasp the ecologically relevance of CIET in coastal sediments and beyond.

### Implications

Long distance electron transfer in subsurface sediments has been proposed to operate as a directional chain of short-distance ET-units^58^. The *Ca*. Geosyntrophus acetoxidans-*Methanosarcina* consortium described here exemplifies one such unit, with electrons flowing via conductive particles between an electrogenic SAO-bacteria and an electrotrophic methanogen, forming a particle-bound CIET module that can be embedded within larger redox-gradient ET networks.

Geoconductors (conductive particles native to soils and sediments) are abundant in the very settings where our mechanisms should matter–anoxic soils and sediments^59–67^. Black carbon (BC), from wildfires, fossil fuel combustion and industrial sources^63,68^ can reach extreme levels near coastlines, glacier bays and river runoffs. For example, BC reached up to 17 mg g⁻¹ dry sediment in heavily polluted Norwegian harbors; ∼8 mg g⁻¹ in some North American estuaries; as much as ∼100% of total organic carbon (TOC) in Arctic glacial bays (e.g., Kongsfjorden) where BC-contaminated particulates are rapidly deposited at glacial fronts; and 46 ± 2% of TOC in Baltic Sea river-outflow sediments—where our consortium is derived from. Beyond BC, iron-bearing geoconductors (magnetite, pyrite, greigite) are widespread: iron is the fourth most abundant crustal element, and reactive, metastable Fe phases (e.g., ferrihydrite, goethite) are enriched in top sediment strata of coastal wetlands. Rivers, atmospheric dust, and hydrothermal inputs continuously deliver these minerals to coastal sediments, while climate-driven erosion^69,70^, glacial melt, wildfires^60,68^, coal combustion^71^, biochar supplementation in agriculture^59^ and other anthropogenic deposition increase geoconductor loads beyond natural levels.

These particle inventories provide persistent niches that may select for EET-enabled metabolisms and interspecies interactions. Such metabolisms are often overlooked in laboratory studies that rely on cells removed from their native matrices. Given the documented geoconductor abundance (black carbon and iron minerals like magnetite) in many coastal and estuary environments, particle-obligate syntrophies are likely hotspots for routing organics to methane. However, distinguishing particle-obligate syntrophy from other methanogenic pathways in situ remains challenging^4^. The molecular signatures identified here (novel PCC-clusters and MmcA variants) provide some of the first genomic markers for targeted detection. It is now critical to (i) survey geoconductor-rich sites (harbours with BC ≳10 mg g⁻¹, polluted estuaries, Arctic glacial bays, Baltic river runoffs) for these markers and CIET-SAO rates; (ii) establish thresholds for particle type, size, and conductivity that sustain CIET; and (iii) obtain in situ rate measurements to quantify the contribution of particle-mediated methanogenesis to methane budgets. Since eroded terrigenous inputs and anthropogenic deposition are increasing, quantifying where and when geoconductors support these alliances is essential for predicting methane emissions under accelerating environmental change.

## Material and Methods

### Cultivation

Over nine years, we conducted more than 22 transfers by adding 10-20% inoculum to fresh DSMZ modified 120 media^15^, which contained 10 mM acetate as the sole electron donor. Control incubations were prepared without conductive particles, while incubations with conductive particles included 10g/L Granular Activated Carbon (GAC, Merck; <5 mm) per culture media.

Before adding the sterile media and inoculum, GAC was wet presterilized. This was done by overlaying it with 200 µL of ultrapure water, then degassing it for 3 minutes with an 80:20 N_2_:CO_2_ gas mix and finally autoclaving it at 121°C for 25 minutes. The media was autoclaved separately and then added to the bottles containing sterile GAC under strict anaerobic conditions^15^.

The serum bottle volume for the inoculations was 100 mL, consisting of 50 mL culture and 50 mL headspace. All subsequent transfers were prepared in 50 mL blue chlorobutyl-rubber-stoppered glass vials. All culture experiments were carried out with at least three biological replicates.

All cultures were incubated in the dark, without shaking, at room temperature (22°C). Subsampling and transfer procedures were performed under strict sterile and anaerobic conditions using gas-tight syringes and hypodermic needles degassed with an 80:20 N_2_:CO_2_ gas mix.

### Analytical measurements

For methane determination, headspace gas samples (1 mL) were withdrawn periodically using sterile, degassed syringes and injected using the gas displacement method in 3 mL gas-tight exetainers (Labco, UK) filled with sterile water (Yee et al., 2019). These methane samples were stored upside down at room temperature until measurement. Headspace methane concentrations were determined by injecting 100 µl in a Trace 1300 gas chromatograph (Thermo-Scientific) equipped with a TG-Bond Msieve 5A column (30 m × 0.53 mm × 50 µm) and a flame ionization detector (FID). Nitrogen was used as the carrier gas at a 5 mL/min flow rate. The oven was maintained at an isothermal temperature of 150°C, while the injector and detector temperatures were set at 200°C. Methane quantification was based on external calibration using in-house prepared gas standards, ranging from 0.01% to 20% CH_4_ in CO_2_. To determine the acetate concentration in the media, we sampled 1 mL of media with a 1 mL gas-tight syringe, filtered through a 0,45 µM filter, and diluted 1:10 with ultrapure water. Acetate samples were then measured on a Dionex ICS-1500 Ion Chromatography System (ICS-1500) equipped with the AS50 autosampler, and an IonPac AS22 column coupled to a conductivity detector (31 mA). The eluent used was 4.5 mM Na_2_CO_3_ with 1.4 mM NaHCO_3_, and the run was isothermic at 30 °C with a flow rate of 1.2 mL/min^26^.

### Helium Ion Microscopy

To investigate the spatial distribution of the microorganisms on the GAC surface, we examined the microorganisms attached to the GAC surface using Scanning Helium Ion Microscopy (HIM). The samples were fixed overnight, in the dark at 4°C, using a solution of 2.5% glutaraldehyde in 0.1 M phosphate buffer (pH 7.3). After fixation, the samples were washed three times in 0.1 M phosphate buffer for 10 minutes each, followed by a dehydration process through a graded series of ethanol (50%, 70%, 80%, 90%, 95%, and 100%) with each concentration being used three times for 10 minutes each^72^. Subsequently, the samples were immersed twice in pure hexamethyldisilazane (HMDS) for 30 seconds and then air-dried for 10 minutes^73^. Imaging was conducted using a Zeiss ORION NanoFAB Helium Ion Microscope with secondary electron (SE) detection (Zeiss, Germany). Helium ion imaging was performed at 30 keV beam energy, with a probe current ranging from 0.129 to 0.152 pA and a scan dwell time of 1 µs. Charge compensation was applied when necessary, using a low-energy electron flood gun (433 eV), and the working distance during imaging ranged from 8.0 to 8.8 mm.

### DNA Extraction and Sequencing

DNA was extracted from a parallel set of GAC incubations when methane production reached mid-exponential (2 mM, at day 42) and from particle-free control incubations on day 42. DNA was purified using the DNeasy PowerLyzer PowerSoil Kit (Qiagen). The extraction process followed a modified manufacturer’s protocol, which included specific centrifugation and resuspension steps.

For bead beating, we used G2 DNA/RNA Enhancer beads 0.1 mm (A420150) for varying durations based on the sample type GAC samples for 7.5 minutes, and GAC-free control samples for 3 minutes. The concentrations and purities of the DNA were measured with the Qubit dsDNA HS Assay kit (Thermo Fisher Scientific, USA) and NanoDrop One (Thermo Fisher Scientific, USA).

DNA size distributions were evaluated using the Genomic DNA ScreenTapes on the Agilent Tapestation 4200 (Agilent, USA). Barcoded SQK-NBD114.96 (AC-C, AC-MGN) and SQK-RPB114.24 (AC-GAC) DNA libraries were prepared according to the manufacturer’s protocol, except for minor modifications (Oxford Nanopore Technologies, Oxford, United Kingdom). The barcoded DNA libraries were loaded onto primed FLO-PRO114M (R10.4.1) flow cells and sequenced on a PromethION P2 Solo device, running MinKNOW Release 23.07.12. Signal data was base called and demultiplexed with Dorado basecall server v. 7.1.4 (Oxford Nanopore Technologies, Oxford, United Kingdom) using the super-accurate algorithm (dna_r10.4.1_e8.2_400bps_5khz_sup.cfg). The remaining adapters were trimmed with Porechop v. 0.2.4. Removal of low-quality reads and generation of basic sequencing data statistics were obtained using Nanoqv. 0.10.0^74^.

### De novo assembly, binning and annotation

De novo assemblies were generated individually for GAC and GAC-free control samples with Flye v. 2.9.2-b1786^75–77^, setting the flags –meta and–extra-params min_read_cov_cutoff=10. The draft assembly was subsequently polished once with Medaka v. 1.8.0 (Oxford Nanopore Technologies, Oxford, United Kingdom). Assembly graphs were inspected with Bandage v. 0.8.1^78^. Contigs below 1000 bp were removed with SeqKit v. 2.2.0^79^. Contigs were subjected to automated binning using MetaBAT2 v. 2.15^80,81^. Intra-assembly MAG filtering and inter-assembly MAG dereplication were conducted using dRep v. 3.4.3, setting minimum MAG length (cluster size) to 0.5 Mbp, minimum completion to 1% and maximum contamination to 10% thresholds^82^. MAG completeness and contamination levels were assessed using CheckM2 v. 1.0.1^83,84^. MAG abundances were calculated using CoverM v. 0.6.1, setting –min-read-percent-identity 95 and –min-read-aligned-percent 90.

The final MAGs were classified against the Genome Taxonomy Database release214 using the Genome Taxonomy Database toolkit v. 2.3.0^85^. Quality filtered DNA sequencing reads were classified with Kaiju v. 1.9.2, against the nr_euk 2023-05-10 database (321 mio. protein sequences from both prokaryotes, microbial eukaryotes and viruses)^86^. Filtered ONT fastq reads were converted to fasta using GNU Awk v. 5.0.1, and genes encoding rRNA were extracted with Barrnap v. 0.9 using a bacterial hidden Markov model. The SILVA 16S/18S rRNA 138 SSURef NR99 full-length database in RESCRIPt format was downloaded from the QIIME on 29 September 2022^87–89^. Potential generic placeholders and dead-end taxonomic entries were cleared from the taxonomy flat file, *i.e.,* entries containing uncultured or unassigned metagenomes, were replaced with a blank entry. Extracted rRNA genes were mapped to the database using Minimap2 v. 2.24-r1122^90^ and sorted using Samtools v.1.15.1^91^. Mapping results were filtered such that alignment sequence length covered > 85% of database entry, and with mapping quality score > 0.85. Further bioinformatic processing was done via RStudio IDE (2022.12.0.353) running R version 4.2.2 (2022-10-31 ucrt) and using the R packages: ampvis2^92^, tidyverse (2.0.0), seqinr, ShortRead and iNEXT^93,94^. Bakta v. 1.8.1^95^ and Prokka v. 1.14.6^96^ were used to annotate the dereplicated bacteria and archaea MAGs using the –compliant flag (Genbank/ENA/DDJB compliance), respectively. The –skip-crispr flag was envoked in Bakta.

### Phylogenomics trees

For phylogenomic analysis of the *Ca*. *Geosyntrophus acetoxidans* and the novel type-II *Methanosarcina*, we used bacterial and archaeal marker gene sets provided by the Genome Taxonomy Database (GTDB; release R214), comprising 120 bacterial marker genes (bac120) and 53 archaeal marker proteins (arch53).

The precomputed alignments for the bac120 dataset were downloaded from the PhyloM.bac120.aln directory, with each marker gene aligned up to ∼1052 bacterial reference genomes. From NCBI, we downloaded amino acid sequences from the type strains of representatives within the *Geobacteraceae* and made our database, including our own *Ca*.

Geosyntrophus acetoxidans and *Geobacter psychrophilus* P39 genomes. Using BLASTp, searches were performed with the following parameters: BLOSUM62 scoring matrix, gap costs of 11 (open) and 1 (extend), E-value threshold of 1e−10, and a maximum of 5 hits per query. The top hit per genome (lowest E-value) was selected for further analysis, and the protein sequences were aligned using MAFFT^97^. Maximum likelihood phylogenetic trees were constructed with RAxML^98^ and 100 bootstrap replicates. Phylogenomic inference was conducted using the PROTCAT model with the DAYHOFF substitution matrix, 25 rate categories, and default seed values (parsimony and bootstrap = 12345). Trees were visualized and annotated using iTOL v6^99^.

The same workflow was applied to archaeal genomes affiliated with the genus *Methanosarcina*. For this, individual alignments of the 53 archaeal marker genes were obtained from the arch53/msa/individual directory of GTDB release R214.

### EET-Machinery Identification

To identify the EET-signatures in our electrogen, we searched for heme-binding motifs in the MAGs obtained from GAC inoculations. We used an in-house script to screen all MAGs protein sequences for the canonical CxxCH motif as well as the extended variants^100^. Each motif was searched separately, allowing for no mismatches. Motif positions were annotated and monitored for each gene to assess the total number and spread of heme-binding sites. To predict the localization of all identified multiheme cytochromes we used PSortB vs. 3.0.3^101^ and DeepLocPro vs.1.0^102^.

Relationship between OMCs in Ca. Geosyntrophus acetoxidans and *Geobacter* OmcB was determined by protein BLAST and homology search to the reference *omcB* sequence from *Geobacter sulfurreducens* strain PCA (RefSeq: WP_286351715.1), which is known to contain 12 heme-binding sites^103^. The sequence was obtained from NCBI to identify homologous proteins within our own *Geobacteraceae* protein database using BLASTp. Top hits from each species were selected and aligned with the MAFFT and a maximum-likelihood tree was constructed with RaxML. To identify the porin of the potential porin–cytochrome–cytochrome (PCC) gene clusters, genomic regions preceding the candidate cytochromes were manually examined. The amino acid sequences of adjacent upstream genes were extracted and submitted to InterPro (release 103.0) for domain analysis. The identification of a porin domain upstream of the multiheme cytochromes served as evidence for the existence of potential PCC structures. The porin domain was validated by the detection of Superfamily SSF56935 (Model: 0048638), covering amino acid positions 259– 387, which align with a conserved outer membrane β-barrel porin domain. These annotations were derived from the InterPro database^104^.

To identify putative electrically conductive e-pili, we utilized the e-pilin protein sequence from *Geobacter sulfurreducens* strain PCA (GenBank accession: AAR34870.1) as a reference^36^. The sequence was aligned against our MAG dataset and a custom *Geobacteraceae* protein database. The same procedure was performed for the archaellum as a reference to the *flaB1* (WP_011023000.1) and *flaB2* (WP_011023001.1) of *M. acetivorans* CA2^37^, following the methodology as mentioned before for omcB.

### Average Nucleotide Identity (ANI) and Average Aminoacid Identity (AAI)

The genome wide average nucleotide identity of *Ca.* Geosyntrophus acetoxidans was calculated against its closest relatives using OAT from EYBiocloud^105,105^. OAT employs the OrthoANI algorithm and BLAST calculations to measure the similarity between two genome sequences. For the calculation of the genome-wide AAI^106^ between closest related proteomes, computed protein sequences were pairwise aligned using DIAMOND v2.0.15. The workflow was computed in our own Python script, and we used the Pandas library for data manipulation and analysis. All the alignments were sorted, prioritizing alignments by query sequence, bit score, and e-value. Here, we calculated the AAI for the alignments, and alignments with a percentage identity of at least 30% and an alignment length of at least 50 amino acids were filtered. From those, only the top-scoring alignment (based on bit score) was retained for each query protein. The AAI was computed as the mean percentage identity across all retained reciprocal best hits (RBHs). This analysis yielded the genome-wide AAI and the number of RBHs, providing a quantitative measure of proteomic similarity between the two genomes.

## Data availability

All metagenome-assembled genomes (MAGs) and annotations are available in the NCBI database under the following accession numbers: Bioproject: PRJNA1140524, Biosamples: SAMN47284751 (MAGs from GAC-incubations), SAMN47284749 (MAGs from particle-free incubations), and SAMN47284752 (whole-genome sequence of *Geobacter psychrophilus* P39).

## Acknowledgments

This work contributes to a Danish Research Council grant DFF Project 2 (grant-number 1026-00159B) awarded to AER. Additional support was provided by MIMET, an ERC Consolidator Grant (grant-number 101045149) to AER. The microscopy imaging was supported by the Interreg 6a Deutschland-Denmark Program project PlastTrack (grant number 07-1-22 1) and by UFM5229-0001B, project NANOCHEM, National Infrastructure awarded to JF.

